# Honeybees exposure to veterinary drugs: how the gut microbiota is affected

**DOI:** 10.1101/2021.03.04.434023

**Authors:** L. Baffoni, D. Alberoni, F. Gaggìa, C. Braglia, C. Stanton, P.R. Ross, D. Di Gioia

**Affiliations:** Department of Agricultural and Food Sciences (DISTAL), University of Bologna, Viale Fanin 44, 40127 Bologna, Italy; Teagasc Food Research Centre, Moorepark, Fermoy, Co. Cork, Ireland; APC Microbiome Institute, University College Cork, Co. Cork, Ireland

## Abstract

Several studies have outlined that a balanced gut microbiota offers metabolic and protective functions supporting honeybee health and performances. The present work contributes to increasing knowledge on the impact on the honeybee gut microbiota of the administration of three different veterinary drugs (oxytetracicline, sulphonamides and tylosin). The trial was designed with a semi-field approach in micro-hives containing about 500 bees, i.e. in experimental conditions as close as possible to real hives considering the restrictions on the use of antibiotics; 6 replicates were considered for each treatment plus the control. The absolute abundance of the major gut microbial taxa in newly eclosed individuals was studied with qPCR and next generation sequencing. Antimicrobial resistance genes for the target antibiotics were also monitored using a qPCR approach. The results showed that none of the veterinary drugs altered the total amount of gut bacteria, but qualitative variations were observed. Tylosin treatment determined a significant decrease of α- and β-diversity indexes and a strong the depletion of the rectum population (lactobacilli and bifidobacteria) while favoring the hindgut population (*Gilliamella*, *Snodgrassella* and *Frischella* spp.). Major changes were also observed in honeybees treated with sulphonamides, with a decrease in *Bartonella* and *Frischella* core taxa an increase of *Bombilactobacillus* spp. and *Snodgrassella* spp. Conversely, minor effects were observed in oxytetracycline treated honeybees. Monitoring of antibiotic resistance genes confirmed that honeybees represent a great reservoir of tetracycline resistance genes. Tetracycline and sulphonamides resistant genes tended to increase in the gut microbiota population upon antibiotic administration.

**Importance:** This study investigates the impact of the three most widely used antibiotics in the beekeeping sector (oxytetracycline, tylosin and sulphonamides) on the honeybee gut microbiota and on the spread of antibiotic resistance genes. The research represents an advancement to the present literature considering that tylosin and sulphonamides effect on the gut microbiota has never been studied. Another original aspect lies in the experimental approach used, as the study looks at the impact of veterinary drugs and feed supplements 24 days after the beginning of the administration, thus exploring perturbations in newly eclosed honeybees, instead of the same treated honeybee generation. Moreover, the study is not performed with cage tests but in micro-hives thus reaching conditions closer to real hives. The study reaches the conclusion that tylosin and sulfonamides determine major changes in some core members and that antibiotic resistance genes for tetracycline and sulphonamides increase upon antibiotic treatment.

## Introduction

Bees have a globally recognized importance for the maintenance of the planet biodiversity and crops pollination (1, 2). In addition, honeybees are valuable for the production of commercially important hive products, such as honey, propolis, royal jelly and wax.

In the last 150 years, farming practices aimed at increasing livestock productivity have been applied and antibiotics have played a crucial role (3) in intensive breeding. Honeybees are not an exception and large-scale apiaries of dozens of beehives have replaced the few hives in the yard of farmers and wild colonies (4). Moreover, hives are often moved for long distances for agriculture pollination needs (5; 6) or shipped worldwide for transnational commercialization (7, 8).

The intensive exploitation of agricultural systems and pollinators resources (9) contribute to honeybee stress in such a way that they can no longer survive without constant anthropogenic inputs (10, 11). Among the biotic and abiotic factors that are affecting honeybee health, pathogens and parasites play the greatest role. These, acting in synergy with abiotic factors, have caused significant decline in the European colonies (12, 13).

In order to fight microbial pathogens, several antibiotics are used, such as oxytetracycline-HCl (Terramicin^®^) against *Paenibacillus larvae* (14), tylosine (Tylovet^®^) against *Melissococcus plutonius* (15, 16), and sulphonamides to control both pathogenic bacteria and, partially, Nosemosis, caused by *Nosema apis* and *Nosema ceranae* (16). The virulence and spread of pathogens are often enhanced by modern beekeeping practices (17), like the unnatural proximity of colonies (18) and frames exchange (19).

Since the early 2000s, concern about the spread of antibiotic resistance genes among pathogenic bacteria has led many nations to apply restrictions on their use on livestock (20, 21). In apiculture, most of the authorizations to trade certain antibiotics have been withdrawn by the European Commission or by pharmaceutical companies themselves (22, 23). Conversely, antibiotic administration to honeybees is permitted, in many other countries, even though with restriction and controls (24, 25), and the European honey market is still threatened by antibiotic residues (26).

The honeybee gut microbiota is relatively simple, composed of few core bacterial genera and other non-core genera with a low or occasional presence (27, 28). Commensal gut bacteria, besides their role in honeybee nutrition and physiology, act in synergy with the host immune system and play a role in modulating the insect response to pathogens (29, 30). The honeybee gut microbiota is directly influenced by various factors such as diet, season and exposure to chemical compounds such as weed killers or antibiotics (31–33) and its unbalance, defined as intestinal dysbiosis (34), may negatively influence honeybee well-being.

In this work, we investigated the effect on the honeybee gut microbial community of the most used veterinary drugs such as oxytetracicline, sulphonamides and tylosin. A number of studies, often based on cage tests or on hybrid approach between cage test and restricted realise time into the hive, have considered the impact of oxytetracycline on the honeybee gut microbiota, whereas, to the best of our knowledge, sulphonamides and tylosin have never been investigated before. This study has been performed using a semi field approach, *e.g.* in experimental conditions as close as possible to real hives taking into account the restrictions on the use of antibiotics, thus partially avoiding artificial conditions typical of the cage tests. Perturbation of the gut microbiota in newly eclosed individuals are explored with the use of qPCR and next generation sequencing (NGS) and antimicrobial resistance genes for the target antibiotics were monitored.

## Results

### General observations on the colonies status pre and post treatment

The trial involved bees treated with tetracycline (PT), sulphonamides (SUL) and tylosin (TL), plus an untreated control (CTR); each experimental condition was tested with 6 replicates. Bees were sampled at T0 (experiment beginning) and T1 (24 days later). Samples are therefore expressed as sampling time – experimental condition – replicate number (e.g.: T0_CTR_1). Moreover, the experiment relied on micro-hives managed with a semi-field approach due to national restriction.

The health status of the treated honeybee micro-hives was generally good all over the trial. Only one micro-hive collapsed (PT_6) just after the experiment end, presumably due to varroosis, whereas CTR_5, PT_1 and SUL_1 were found to be queenless during the experiment. Visual evaluation at the time of gut sampling highlighted a reddish coloration of the intestinal epithelium in the tylosin treatment group. Drought conditions in the second half of the experiment did not allow nectar harvest.

### qPCR quantification of total bacteria, *Bifidobacterium* and Lactobacillaceae in the gut

The count of Eubacteria (Fig. 1A) at the beginning and at the end of the experiment showed a significant decrease (0.65 Log, p<0.05) upon sulphonamide treatment (SUL_T0 *vs* SUL_T1); the other treatment did not show significant variation. A non-significant decreased was observed in the control micro-hives (CTR_T0 *vs* CTR_T1) with a loss of 0.21 Log 16S rRNA copies/intestine. *Bifidobacterium* spp. counts showed a general decrease in all experimental conditions. The reduction was significant in PT_T0 *vs* PT_T1 (0.58 Log CFU/intestine, p<0.01) and in TL_T0 *vs* TL_T1 (3.61 Log CFU/intestine decrease, p<0.01; Fig. 1B). Also, Lactobacillaceae showed a general decrease in all experimental conditions, which was significant only in the comparison of TL_T0 *vs* TL_T1 (p<0.01) with a decrease of 0.56 Log CFU/intestine (Fig. 1C).

**Fig. 1A-1C.**
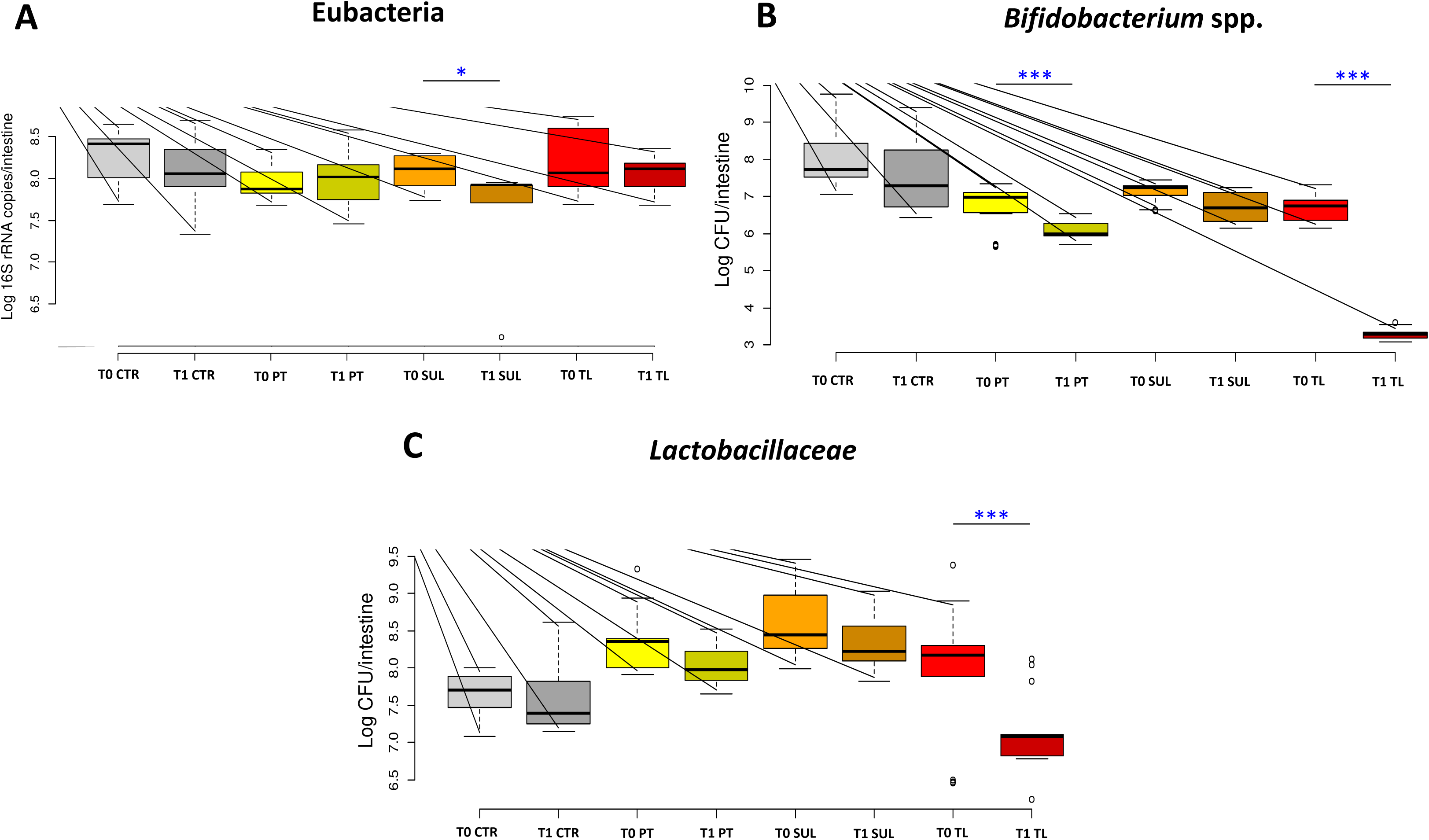
qPCR. quantification of (A) total bacteria (Eubacteria), (B) *Bifidobacterium* spp. and (C) *Lactobacillaceae*. Data are expressed in Log CFU/intestine for *Bifidobacterium* spp. and Lactobacillaceae; for Eubacteria data are expressed as Log 16S rRNA copies/intestine. Boxplots report minimum and maximum values, lower and upper quartile and median. **Antibiotics:** [CTR] Antibiotics Control, [PT] oxytetracycline, [SUL] sulphonamides, [TL] tylosin.

### Bee Gut microbiota analysis via NGS

A total of 48 samples [2 sampling times, T0 and T1, 4 experimental conditions (CTR, PT, SUL, TL), 6 replicates for each condition, each replicate being a pool of 30 honeybee guts] were subjected to NGS analysis on Illumina MiSeq platform. About 13.7 million raw reads were obtained from the sequencing. 9.1 million reads passed the quality control and the Chimera check analysis obtaining an average of 95,986 joint reads per sample. For statistical analysis, samples were rarefied at 48,400 reads, a value obtained excluding one replicate (TL_T1_4) due to a particularly low coverage. The taxonomical assignment of the 47 samples produced 17,194 OTUs at 97% similarity based on SILVA 132 database. The elaboration of NGS data on the whole dataset is reported in Table 3, where absolute abundance at phyla, family and genus level are reported per treatment and time, whereas Fig. 2A reports absolute abundance at genus level per replicate.

**Table 1.**
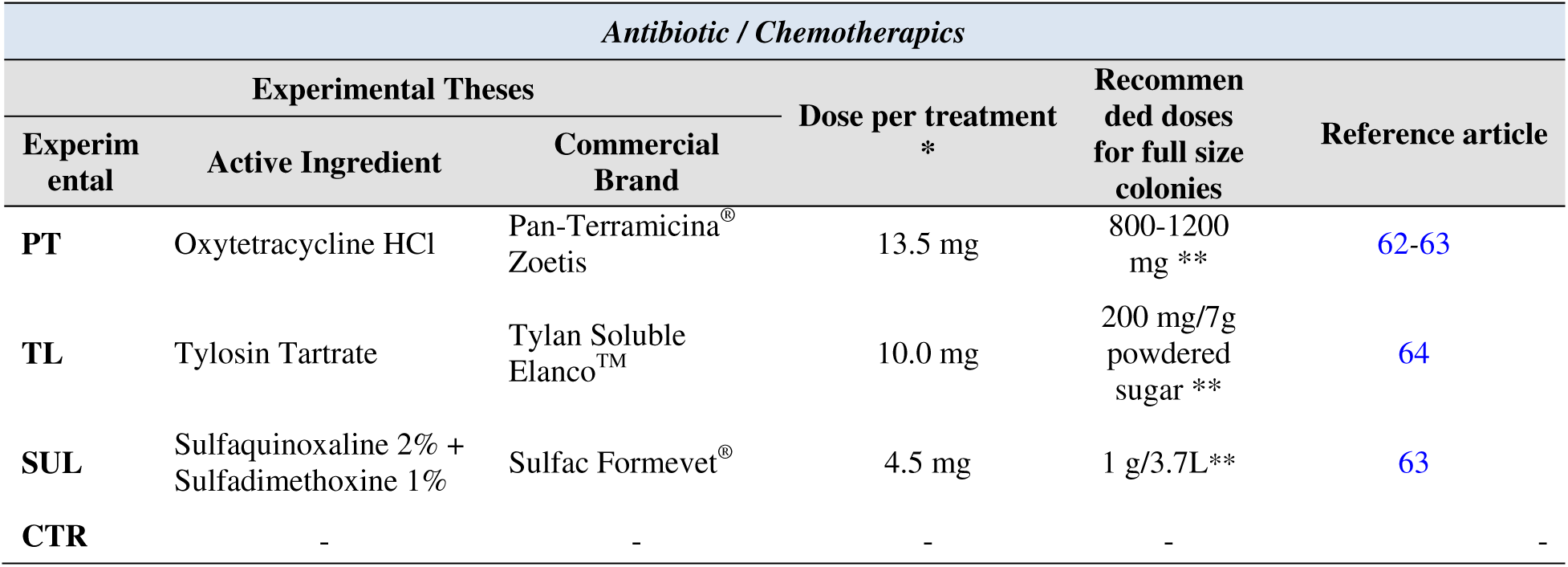
Antibiotics used in this work, their dosages applied in each treatment per hive in the presented trials, and recommended doses for full size colonies. All antibiotics or antimicrobial agents were prepared in 30 mL of sugar syrup and sprayed on, or fed to bees. *Dose recalculated according to the colony size of microhives, expressed as mg or μL of active ingredient dissolved in 30 mL of sugar syrup. **Total recommended dose for 3 administrations with weekly cadence;

**Table 2.**
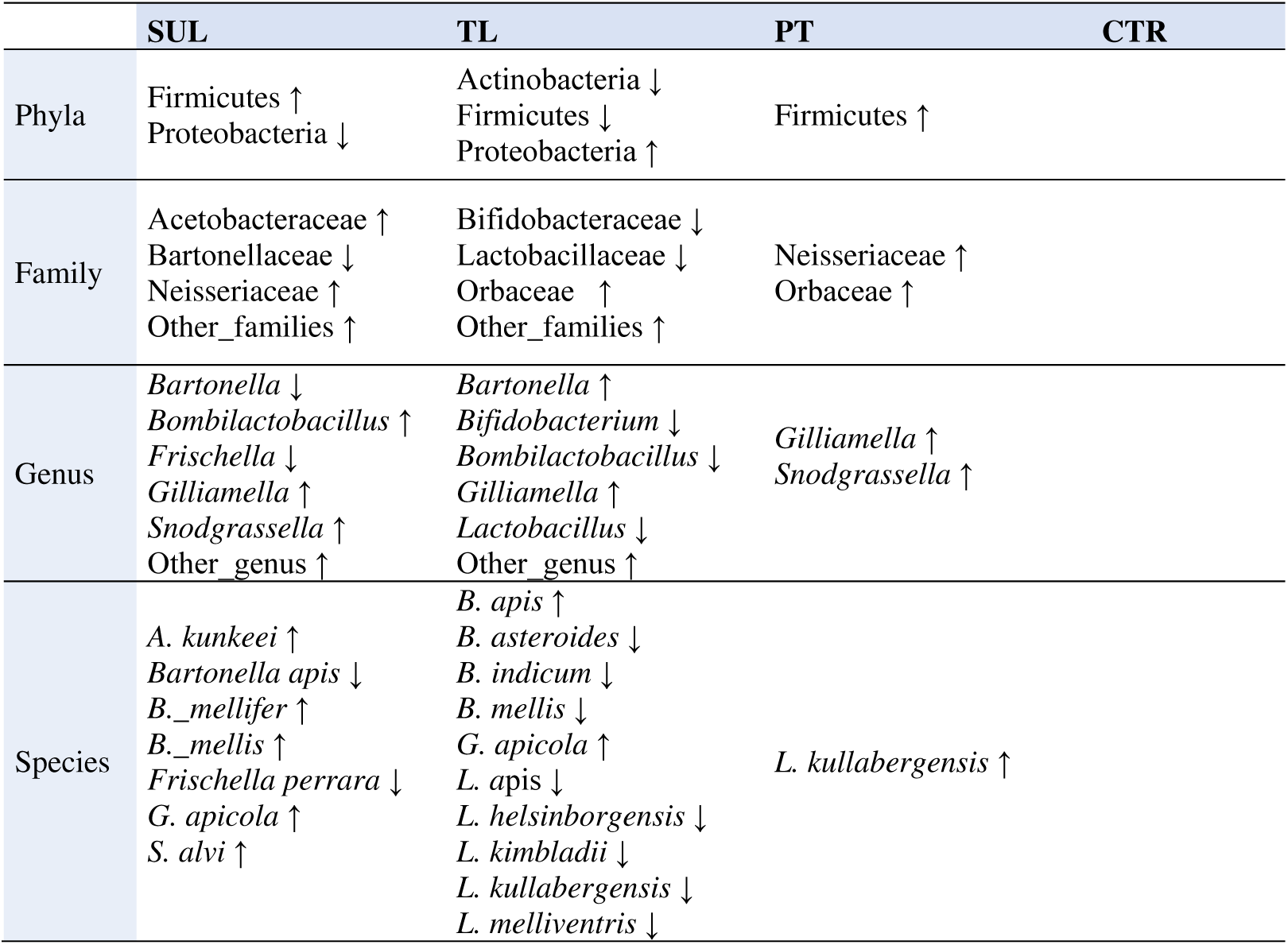
Significant variation among microbial groups at phyla, family, genus and species level according to the experimental conditions.

**Table 3.**
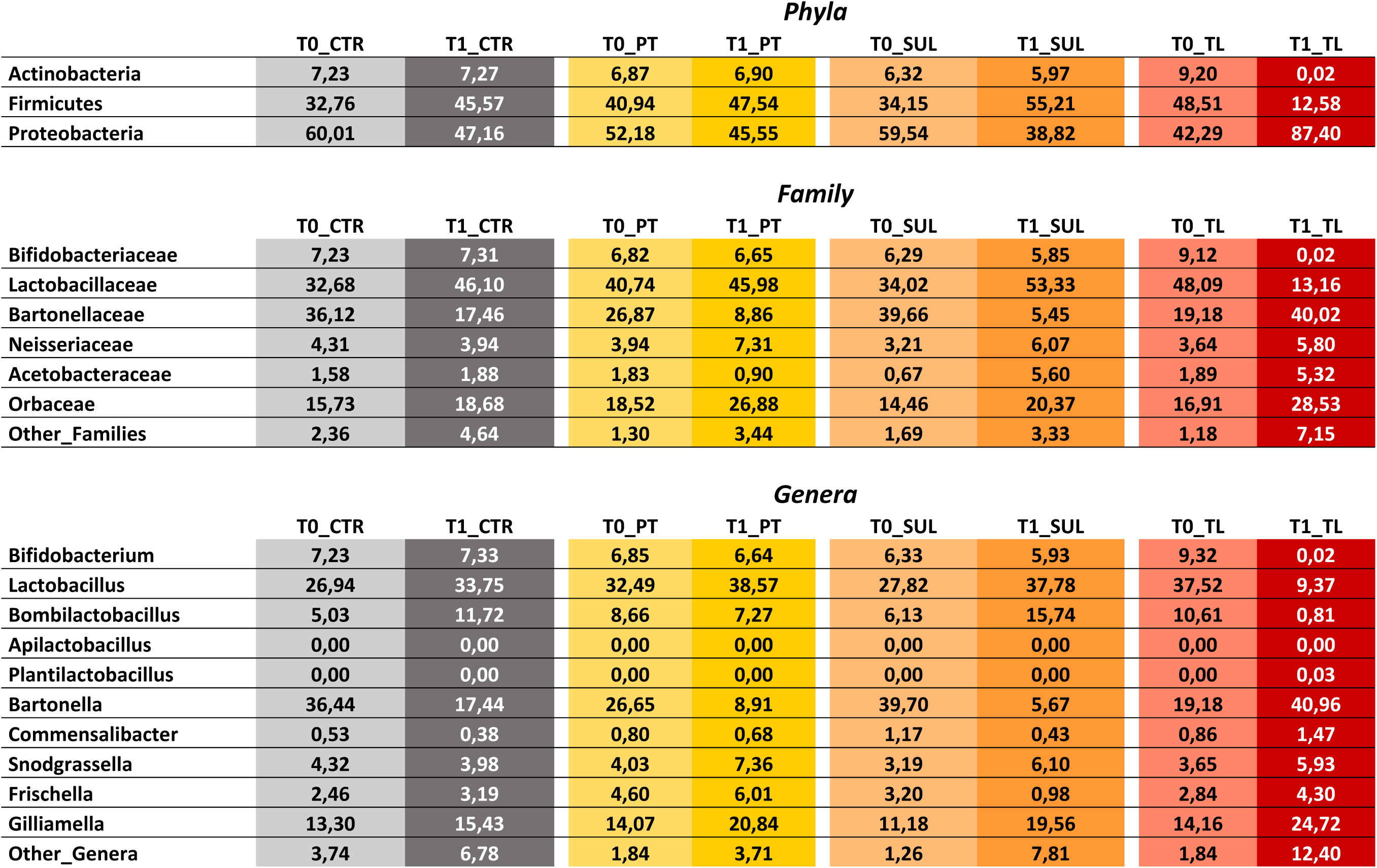
NGS absolute abundance at phyla, family and genus level, reported per treatment and sampling time.

**Fig. 2A-2C.**
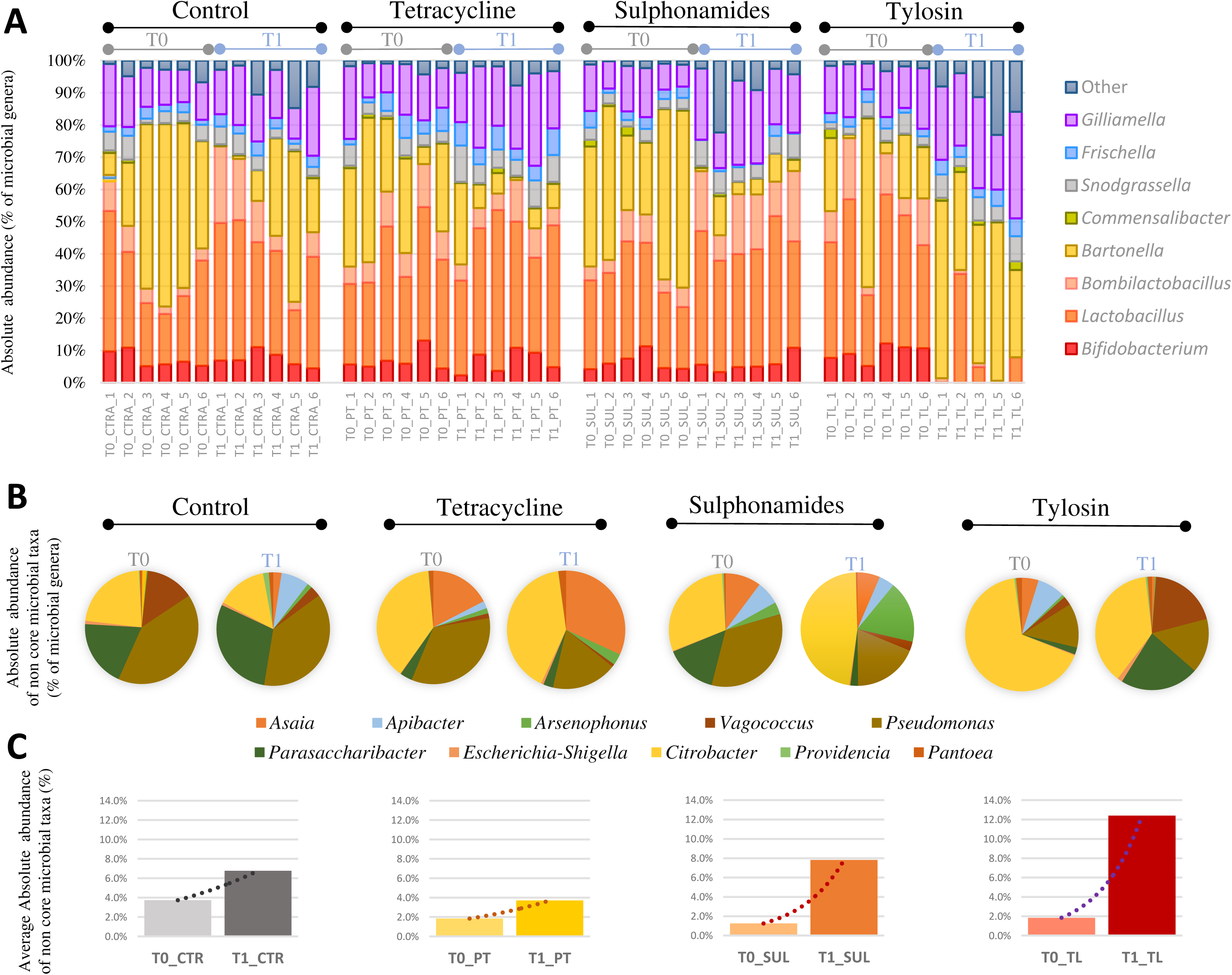
NGS Absolute Abundance overview. (A) bar charts reporting the major cumulated microbial genera per samples and their absolute abundance expressed in percentage. (B) pie-charts reporting the minor cumulated microbial genera (Other_taxa) per experimental conditions and sampling time, expressed in percentage as absolute abundance. (C) average absolute abundance of Other_taxa for each treatment in T0 and T1.

Detected non-core genus could be mainly ascribed to the genera: *Asaia*, *Apibacter*, *Arsenophonus*, *Vagococcus*, *Pseudomonas*, *Parasaccharibacter*, *Citrobacter*, *Providencia* and *Pantoea* (Fig. 2B) and their proportion at T0 and T1 (Fig. 2C).

α-diversity indexes (Chao1, Observed OTU and PD whole tree) showed a significant decrease over time only in tylosin treated group (p<0.01; Fig. S1). The analysis of β-diversity (Table S1) underlined statistically significant differences in the unweighted UniFrac analysis, which stresses the importance of taxa presence/absence, only comparing CTR to TL treatment. However, considering the abundance of taxa in the weighted UniFrac, not only TL treatment resulted significant but also SUL when compared to CTR (Table S1).

### Antibiotic effect

Control bees did not show any significant shift of the intestinal microbial taxa at the different taxonomic levels, comparing the two sampling times. A summary of the significant changes from phyla to species for each antibiotic treatment between the two sampling times level is reported in Table 2.

PT treatment, at phylum level, showed an increase of Firmicutes (from 40.9% at T0 to 47.5% at T1) and a decrease of Proteobacteria (from 52.2% to 45.6%), although not significant, whereas Actinobacteria remained stable. At family level, comparing PT_T1 *vs* PT_T0, Bartonellaceae showed a decreasing trend but not significant (from 8.66% to 7.27%), while both Neisseriaceae and Orbaceae significantly increased from 3.94% to 7.31% (p<0.01) and from 18.5% to 26.7% (p<0.05), respectively. At genus level, *Gilliamella* spp. almost doubled its absolute abundance comparing PT_T1 *vs* PT_T0 (from 14.07% to 20.84%; p<0.05), while *Snodgrassella* significantly increased (from 4.03 to 7.36; p=0.01; Fig. 3D). At species level, PT treatment determined a significant increase only for *Lactobacillus kullabergensis* (p<0.01).

**Fig. 3A-3F.**
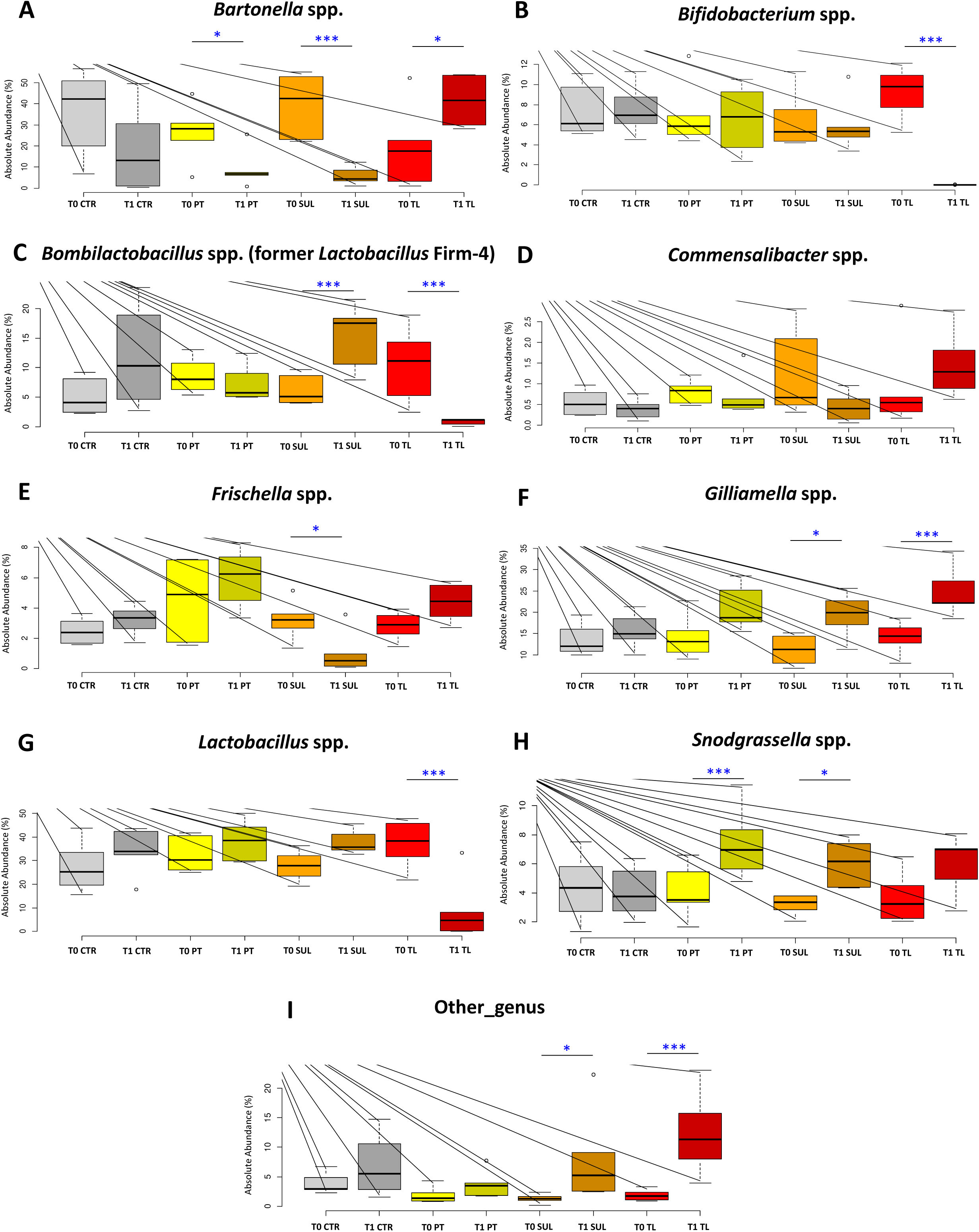
NGS Absolute Abundance at genus level. Box plots reporting the major microbial genera expressed for their absolute abundance in percentage, and in relation to experimental conditions (significant pairwise comparisons *p < 0.05; ***p < 0.01). Boxplots report minimum and maximum values, lower and upper quartile and median. **Microbial taxa described:** (A) *Bartonella* spp., (B) *Bifidobacterium* spp., (C) *Bombilactobacillus* spp., (D) *Commensalibacter* spp., (E) *Frischella* spp., (F) *Gilliamella* spp., (G) *Lactobacillus* spp., (H) *Snodgrassella* spp., (I) Other_genus, for the experimental conditions: [CTR] Control, [PT] oxytetracycline, [SUL] sulphonamides, [TL] tylosin.

Tetracycline resistance gene *tetB* increased significantly of 159% (p<0.01) comparing PT_T1 *vs* PT_T0. However, the increase was also significant comparing CTR_T1 *vs* CTR_T0 (p<0.01). Also, *tetY* drastically increased comparing PT_T1 *vs* PT_T0 (p<0.01) whereas CTR did not show any significant changes.

Regarding SUL treatment, at phylum level, Firmicutes showed a significant increase comparing SUL_T1 *vs* SUL_T0, from 34.1% to 55.2% (p<0.05). On the contrary, Proteobacteria decreased significantly from 59.5% to 38.8% (p<0.05). Actinobacteria slightly decreased from 6.31% at T0 to 5.97% at T1 although not significantly. At family level, Bartonellaceae decreased after treatment (from 39.66% to 5.45%; p<0.01) (Fig. 3C), while Neisseriaceae and Acetobacteraceae significantly increased from 3.21% and 0.67% at T0 to 6.07% and 5.60% at T1, respectively (p<0.05).

At genus level, SUL treatment at T1 determined a significant decrease in the absolute abundance of *Bartonella* spp. reflecting the proportions reported at family level (p<0.01; Fig. 3A), and *Frischella* spp. (from 3.20% to 0.98% p<0.05; Fig. 3E). On the other hand, absolute abundance increased in *Bombilactobacillus* spp. (from 6.13% to 15.74%; p<0.01; Fig. 3C), *Gilliamella* spp. (from 11.18% to 19.56%; p<0.05; Fig. 3F), *Snodgrassella* spp. (from 3.19% to 6.10%; p<0.05; Fig. 3H) and Other_genus (p<0.05; Fig. 3I). At species level, a significant increase was reported for *A. kunkeei* (p>0.05), *Bombilactobacillus mellifer* (p<0.01) and *Bombilactobacillus mellis* (p<0.01). *Bartonella apis*, *Frischella perrara* and *Gilliamella apicola* reflected the genus trend, being the only species within the respective genus.

Sulphonamides resistance gene *sul1* and *sul2* showed a significant increase of 76.84% and 33.95%, respectively, comparing SUL_T1 *vs* SUL_T0 (p<0.01) respectively, whereas *sul3* did not produce any amplification at the different annealing temperatures tested (40, 44, 48, 52, 56, 60 and 64 °C).

Proteobacteria doubled their abundance comparing TL_T1 *vs* TL_T0, from 42.3% at T0 to 87.4% at T1 (p<0.01). On the other hand, both Firmicutes and Actinobacteria significantly decreased from 48.5% to 12.6% (p<0.01) and from 9.19% to 0.024% (p<0.01), respectively. Bifidobacteriaceae and Lactobacillaceae significantly decreased between TL_T1 and TL_T0 (p<0.01) with percentage values that are consistent with those reported below at the genus level. Orbaceae significantly increased from 16.9% at T0 to 28.5% at T1 (+68.63%, p<0.01). Finally, absolute abundance of Other_families significantly increased after TL treatment, from 1.18% at T0 to 7.15% at T1 (+673%, p<0.01). *Bifidobacterium* spp. absolute abundance reduction after TL treatment was highly significant (P<0.01), decreasing from 9.32% at T0 to 0.02% at T1 (Fig. 3B). In the same way, *Bombilactobacillus* spp. and *Lactobacillus* spp. decreased from 10.61% and 37.52% at T0 to 0.81% and 9.37% at T1 (p<0.01; Fig. 3C and 3G), respectively. Moreover, *Bartonella* spp. doubled the absolute abundance (from 19.18% to 40.96%; p<0.05; Fig. 3A) together with *Gilliamella* spp. and Other_genus in TL_T1, that significantly increased from 14.16% and 1.84% at T0 to 24.90% and 12.40% at T1, respectively (p<0.01; Fig. 3F and 3I). At species level, a significant decrease of six *Lactobacillus* species and also of unclassified *Lactobacillus* spp. was observed (p<0.01), together with the decrease of *B. mellis* (p<0.01), *B. asteroides* (p<0.01) and *B. indicum* (p<0.05). The Cramer V test showed a strong biological relevance in pairwise comparisons of TL_T1 *vs* TL _T0 and SUL_T1 *vs* SUL _T0 (Cramer V = 0.53 and 0.45 respectively) (35). PT_T1 *vs* PT _T0 and CTR_T1 *vs* CTR _T0 biological relevance was moderate (Cramer V = 0.25 and 0.23) but not negligible. Tylosin resistance gene *tlrB* and *tlrD* did not showed any significant variation in normalized data.

PCA analysis of the dataset at species level PC1 and PC2 together explain only 25% of the variability. However, TL_T1 group is clearly separated from TL_T0 and also from the other treated samples at T1 (Fig. 4A), particularly along the PC1. Orbaceae and thus *Gilliamella* spp. are associated with TL_T1 as also confirmed by statistical analysis (Fig. 4B and 4C). The graph also shows a clear separation of SUL_T0 and T1 along PC2.

**Fig. 4A–4F.**
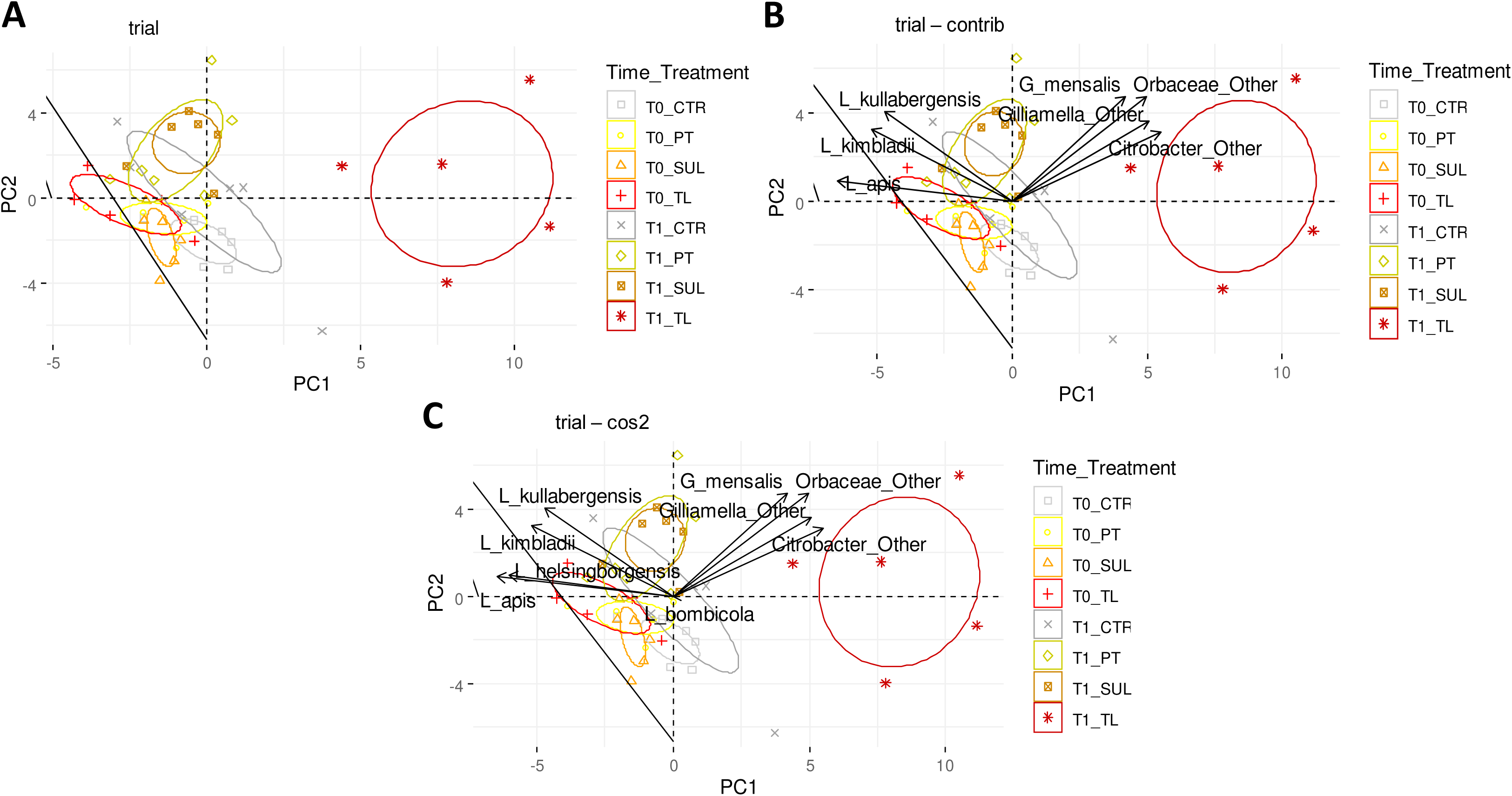
PCA analysis. (A) PCA was performed with 71 taxa at species level, confidence ellipses are shown in the graphs. (B) The graph includes the top seven variables with the highest contrib. (C) The graph includes the variables with cos2>0.6.

**Fig. 5A-5G.**
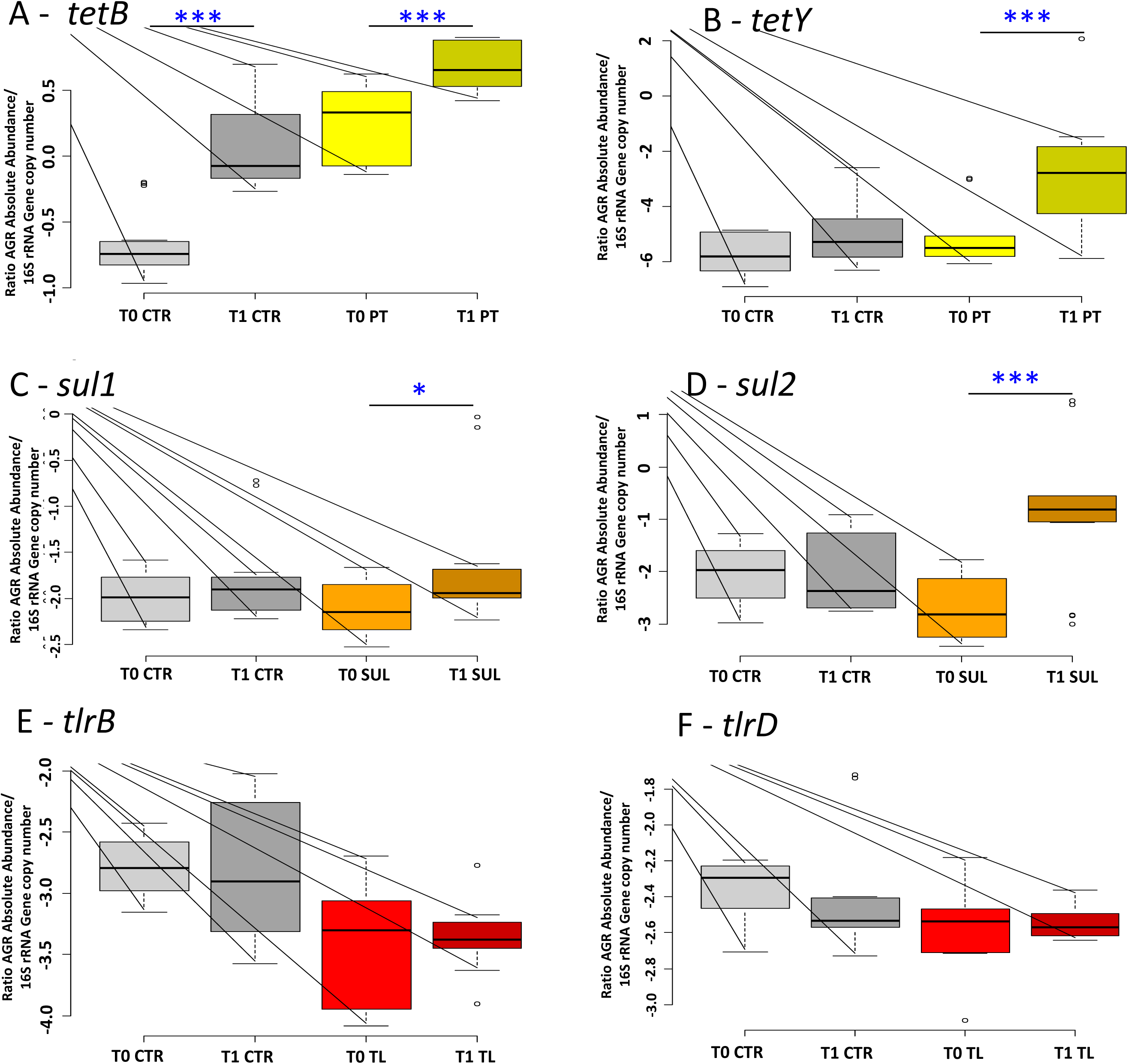
Antibiotic resistance gene: Box plots reporting the AGRs for (A) *tetB*, (B) *tetY* for tetracycline resistance genes; (C) *sul1* and (D) *sul2* for sulphonamides resistance genes; (E) *tlrB* and (F) *tlrD* for tylosin resistance genes. The absolute AGR quantification is normalized with the total 16S rRNA gene copies, in relation to experimental conditions (significant pairwise comparisons *p < 0.05; ***p < 0.01).

## Discussion

This work investigates the gut microbial community of honeybees, which have not been treated with antibiotics for several generations after the supplementation of antibiotics (oxytetracycline, sulphonamides and tylosin).

The observed decrease of total bacterial in treated and control bees could not be ascribed to the antibiotic treatment, but, rather, it seemed to be related with the bee physiology, or stress due to the limited freedom. However, the antibiotic exposure significantly influenced some gut microbial groups.

Oxytetracycline is a broad-spectrum antibiotic currently used in the beekeeping sector (24, 36). Recently, Raymann *et al.* (31, 37) showed that the use of tetracycline strongly decreased the absolute abundance of 5 gut core genera in partially caged honeybees, with a significant decrease of *Bartonella*, *Bifidobacterium*, *Bombilactobacillus* spp. (formerly known as *Lactobacillus* Firm-4), *Lactobacillus* and *Snodgrassella.* Our findings suggest a possible resilience mechanism to the disturbance imposed by oxytetracycline since variations were observed only in two core members (*Bartonella* and *Snodgrassella*) and no significant changes were found in the studied diversity indexes. It is ascertaining that honeybee gut commensal bacteria provide large reservoirs of tetracycline resistance determinants (*otr* and *tet* genes) frequently acquired through massive and/or long-term antibiotic exposure or from other ecosystems shared with animals and humans (38, 39). Ludvigsen et al. (39) showed that honeybee gut symbionts, in particular *Snodgrassella* spp. and *Gilliamella* spp., can survive and proliferate thanks to *tet* determinants. Recently, Daisley *et al.* (40) found that the routine administration of oxytetracycline increases *tetB* and *tetY* abundance in the gut microbiota of adult workers associated with a depletion of the major symbiont taxa. The present study therefore confirms that honeybees represent an impressive reservoir of tetracycline resistance genes, even after two decades without antibiotic treatment. As already mentioned, our experiments were performed on the new honeybee generation, differently from other studies that targeted bees of the same generation (37-38; 40). Beside antibiotic resistant genes uptake, bees, with their daily activities (hive interaction, flying, flower visiting), have a preferred path to replenish their gut microbiota. Most of the published studies rely on caged or partially caged honeybees, which limits social behavior, interactions with the environment but also honeybees queen and brood pheromones for social regulation and interactions. In addition, our work was performed in micro-hives and, therefore, the reservoir of microbial inoculants present in the hive structure (stored pollen, nectar and wax foundation) may have contributed to the mitigation of tetracycline impact.

Sulphonamides (SUL) have been widely used in the beekeeping sector from 1960 to 2000, but residues in honey are still found, thus showing that they are still used in spite of the banning (41). Among the core genera found in the honeybee gut, *Frischella* and *Bartonella* spp. were significantly affected by SUL treatment, while *Bombilactobacillus* spp. and *Snodgrassella* spp. increased their counts. *Frischella perrara* has implications in immune priming in honeybees and in the induction of peptides with antimicrobial activity (42). The registered 3% reduction (with a final 1% abundance in T1) could be detrimental for the bee defense mechanisms. *Bartonella* spp. has been related to the recycling of nitrogenous waste products into amino acids and with the degradation of secondary plant metabolites. The reduction of more than 80% of this taxon could have implication in digestion functions and in the recovery of amino acids (43). However, it is evident that most of the core members are not affected by SUL treatment. This can be again a consequence of the uptake of sulphonamides resistance genes, that was confirmed with both gene *sul1* and *sul2* in this research. This is coherent with results recently obtained by Cenci-Goga *et al.* (44) that found sulphonamide resistance genes (*sul1* and *sul2*) in a large number of honeybees sampled in different Italian locations. Tylosin induced a remarkable change in some microbial taxa proportion, almost causing the depletion of the rectum population (lactobacilli and bifidobacteria) and favoring the hindgut population (mostly *Gilliamella*, but also *Snodgrassella* and *Frischella*). It is known that tylosin targets are mainly Gram-positive bacteria (45; 46). Both *Bifidobacterium*, *Bombilactobacillus* and *Lactobacillus* genera represented 99.99% of Bifidobacteriaceae and Lactobacillaceae family members that, overall, accounted for more than a half of the honeybee gut microbial community. They play an essential role in the transformation of various pollen coat-derived compounds, including flavonoids, phenolamides and ω-hydroxy acids (47), in addition to the complex sugars’ breakdown (48, 49). Their rapid decrease may affect honeybee ability to metabolize specific compounds and consequently reduce nutrient availability. It is remarkable that macrolide antibiotic resistance genes *tlrB* and *tlrD* did not increase significantly in treated honeybees at T1, even if detected. This is probably due to the low occurrence of these antibiotic resistance genes (ARG) in *Bombilactobacillus*, *Lactobacillus* and *Bifidobacterium* honeybee strains, even if TL resistant strains are described in humans and swine (50, 51). *Tlr* genes belong to the same resistance group of *erm* genes (erythromycin ribosome methylation), so that *tlrB* is also classified as *erm32* whereas *tlrD* as *ermN* (52, 53). The presence of *tlr* genes and the lack of decrease upon TL treatment may also be explained by their activity against other macrolide antibiotics that have a broader spectrum of activity, including Gram-negative bacteria that survived the TL treatment. Jackson et al. (54) found that erm genes can be activated after tylosin use. Vice versa *tlr* genes might confer resistance to some macrolide in tylosin unsensitive Gram-negative bacteria populating the honeybee gut thus explaining their presence at T0 and T1.

Several studies showed that environmental species, such as members of the *Asaia, Apibacter, Apilactobacillus, Vagococcus, Pseudomonas, Parasaccharibacter, Citrobacter, Providencia* and *Pantoea* genera, often related with soil, pollen and nectar (55, 56), are detected in the honeybee gut as minor groups (57–59). These non-core genera were found to increase at T1 upon treatments with SUL and TL. These microorganisms may increase the pool of ARG, due to their continuous exposure to antibiotics used in agroecosystem (e.g.: sewage from livestock distributed on soil). For instance, *Parasaccharibacter apium*, recently reclassified as *Bombella* sp. by Smith et al., (60), is reported as a strong immune stimulating strain in honeybees, also capable of counteracting *Nosema* sp. (61). The non-core genera that are sporadically associated with honeybees might play a role in the immune stimulation or metabolic regulation of honeybees, despite their low abundance. Interestingly, the limited interaction with the environment did not prevent their acquisition as gut commensal bacteria over the experimental time.

Overall, the three assayed veterinary drugs do not impact quantitatively the gut bacterial community in terms of total amount of bacteria, but they influence the absolute abundance of several core taxa, causing a possible lack of metabolic functions related to the most susceptible bacterial species and strains. A long-term observation of the colony health status, also including the hive development and hive products (*e.g.* honey), will allow the understanding of the relationship between the altered microbial structure and the behaviour and performance of honeybees.

## Experimental Procedure

### Experimental design

Due to the European and national law restricting the use of antibiotics or other veterinary drugs as antimicrobial in open field, these were tested in semi-field conditions, i.e. in micro-hives incubated in a thermostatic chamber with a limited flying time for honeybees. Honeybees employed in this study have not been treated with antibiotics for several generations (over two decades).

The micro-hives employed in the study were obtained as depicted in Fig. 6A. Shook swarming of a fully populated and healthy bee hive was used to populate 72 micro-combs (L 9.5 x H 10.5 cm). The queen was allowed lying eggs for three days on approximately 1/3 of the total available micro-combs. 5 days later, 24 experimental wooden micro hives (L 20 x H 15 x W 16 cm) were set up, each containing 3 micro-combs (a brood frame, a honey frame and an empty comb). Each micro hive contained approximately 500 honeybees with a mated queen. The obtained micro-hives constituted the experimental replicates (6 for each experimental condition). Moreover, every micro hive was equipped with an anti-robbing entrance modification, forcing honeybees to walk a “S” path that discouraged the entrance of robber bees when the micro hives were placed outside.

**Fig. 6.**
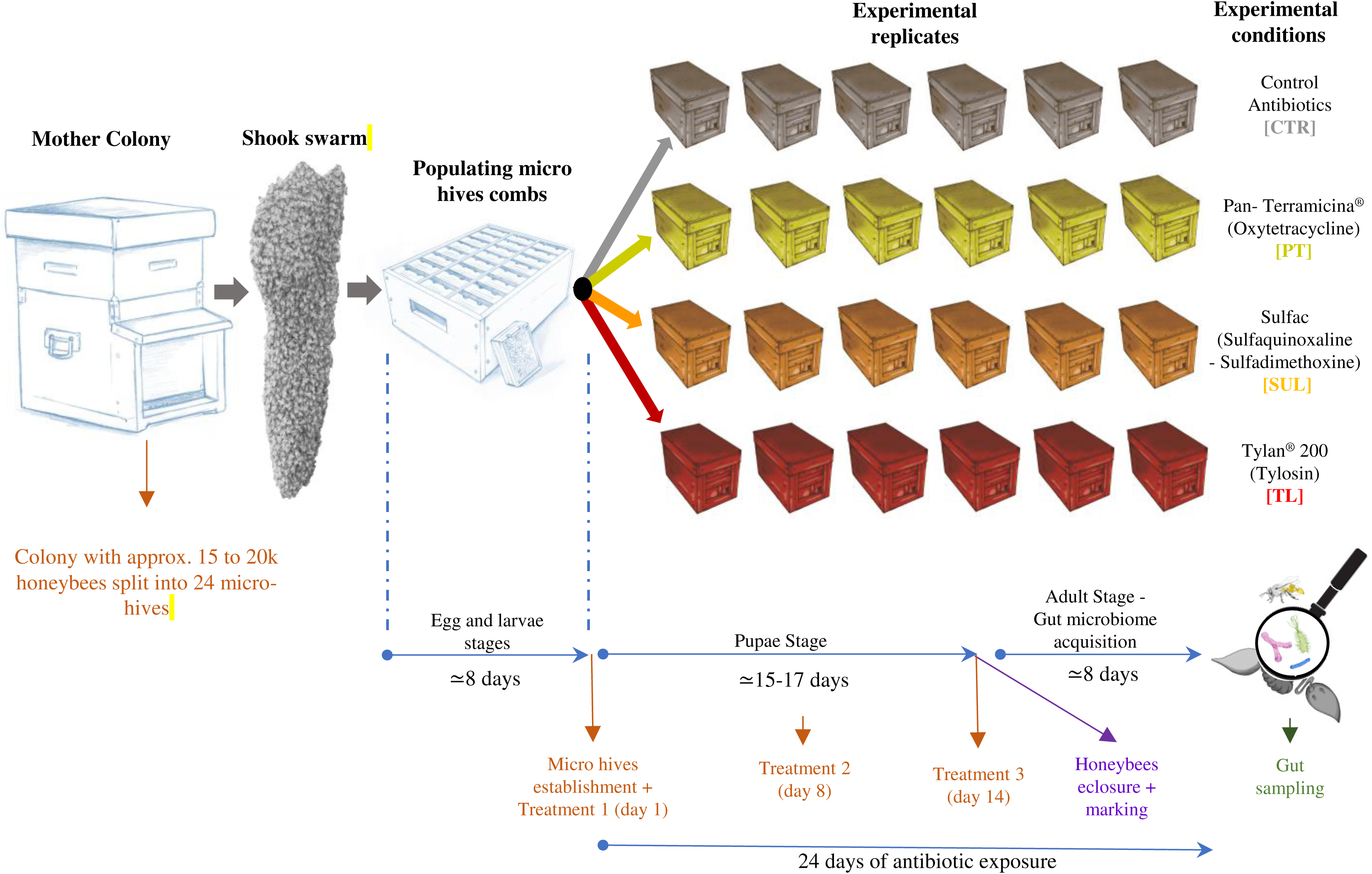
Experimental Design. The figure reports the scheme of the tests and the number of bees and beehives used in the trials.

Micro-hives were located into an incubator with controlled temperature and humidity (29°C and 60 RH), and well equipped with a net allowing ventilation on the mini-hive bottom. The micro-hives were moved outside in the late afternoon (approx. from 5.30 pm to 8.30 pm) every second day in order to allow the bees to fly freely and defecate. The arrangement of the micro hives outdoor in the experimental field always followed the same pattern to avoid disorientation and drift. Micro hives were placed at minimum 2 m distance from each other, and in clusters of 3 units of the same experimental thesis, oriented in different directions, in an experimental forest well populated by trees. At early night-time, micro-hives were closed and re-allocated in the lab thermostat. Micro hives were fed every two days with a 30 ml 1:1 (*w:w*) sucrose solution, plus a 5 ml sterile water dispenser. The day of the antimicrobial treatment, honeybees were treated as described below. The developed experimental conditions were: [TL] tylosin, [PT] oxytetracycline, [SUL] a mixture of sulfaquinoxaline and sulfadimethoxine, and the control with no antibiotic administration [CTR]. Details on antibiotics use and concentration are reported below.

The trial was carried out between July and August, 2016, where two foraging options were available: *Metcalfa pruinosa* honeydew in the early august and *Medicago sativa* blooming all along the trial even if strongly limited by summer drought. The health status (adult honeybee population and brood size, honey reserves, core colony cohesion, symptoms of viral diseases and varroa infestation) of honeybee micro hives was periodically assessed, and variations annotated when relevant.

### Treatment preparation, administration and sampling

Antibiotics were administered according to available guideline for each antibiotic (62–64). Details and concentrations of antibiotics are reported in Table 1. Bees were treated once a week for three weeks with micro hive feeders containing 30 mL of sugar syrup (1:1 w/w) mixed with the respective treatment. Finally, in the days after the 3^rd^ treatment (days 15-17), at least 50 emerging honeybees per replicate were marked on the thorax (65) with coloured nail polish non-toxic to bees. Marked honeybees were sacrificed at day 24, at nurse stage (7-9 days post eclosure) and with a completely established gut microbiota (66). A pool of 30 bees per replicate (a total of 180 samples/experimental condition) was picked at the beginning of the experiment (T0) and after 24 days (T1).

### DNA extraction and NGS sequencing

Obtained honeybee gut pools were well homogenised with pestles, with addition of 1400 µl lysis solution improved with 60 µl proteinase K per pool (20 mg/ml concentration), and glass beads until total destruction of gut epithelial tissues after 1-hour incubation at 55°C. Only 1/4 of the resulting sludge (450 µl) was used for gut genomic DNA extraction with Quick-DNA Fecal and Soil Microbe Kit (Zymo Research, California, U.S.A). The 16S rRNA gene amplification and libraries preparation for Illumina MiSeq platform sequencing were carried out according to Alberoni et al., (67). Bioinformatic analyses were performed with Qiime1, and representative OTUs blasted against the most updated SILVA database release 132. The database was implemented inserting full length 16S rRNA sequences of administered beneficial bacteria. OTUs with less than 0.1% abundance were discarded. α–diversity was evaluated using Chao1, Observed OTU and PD whole tree metrics, whereas β–diversity was evaluated using both weighted and unweighted UniFrac.

### Quantification of target microbial groups and resistance genes

Total bacteria (Eubacteria), Lactobacillaceae family, *Lactobacillus* spp., *Bombilactobacillus* spp. and *Bifidobacterium* spp. were quantified with qPCR (StepOne™ Real-Time PCR System, Applied Biosystems) according to Baffoni *et al.,* (68–69). Data for Lactobacillaceae (*Apilactobacillus* spp., *Bombilactobacillus* spp., *Lactobacillus* spp. and *Lactoplantibacillus* spp.) and *Bifidobacterium* spp. were transformed to obtain the number of microorganism as Log CFU/single intestinal content (70, 71). For total bacteria data were expressed as Log 16S rRNA copies/intestine (72). ARG genes *TetB, TetY, Sul1*, *Sul2*, *Sul3*, *TlrB* and *TlrD* were quantified according Zhang et al. (73). Primers used are reported in Table 4. Raw data were corrected according to the total DNA quantification. The final absolute abundance of ARG was normalized according to (82, 83) by dividing the total ARG with the absolute abundance of total bacteria previously obtained, data reported show the ratio between ARG and total bacteria.

**Table 4.**
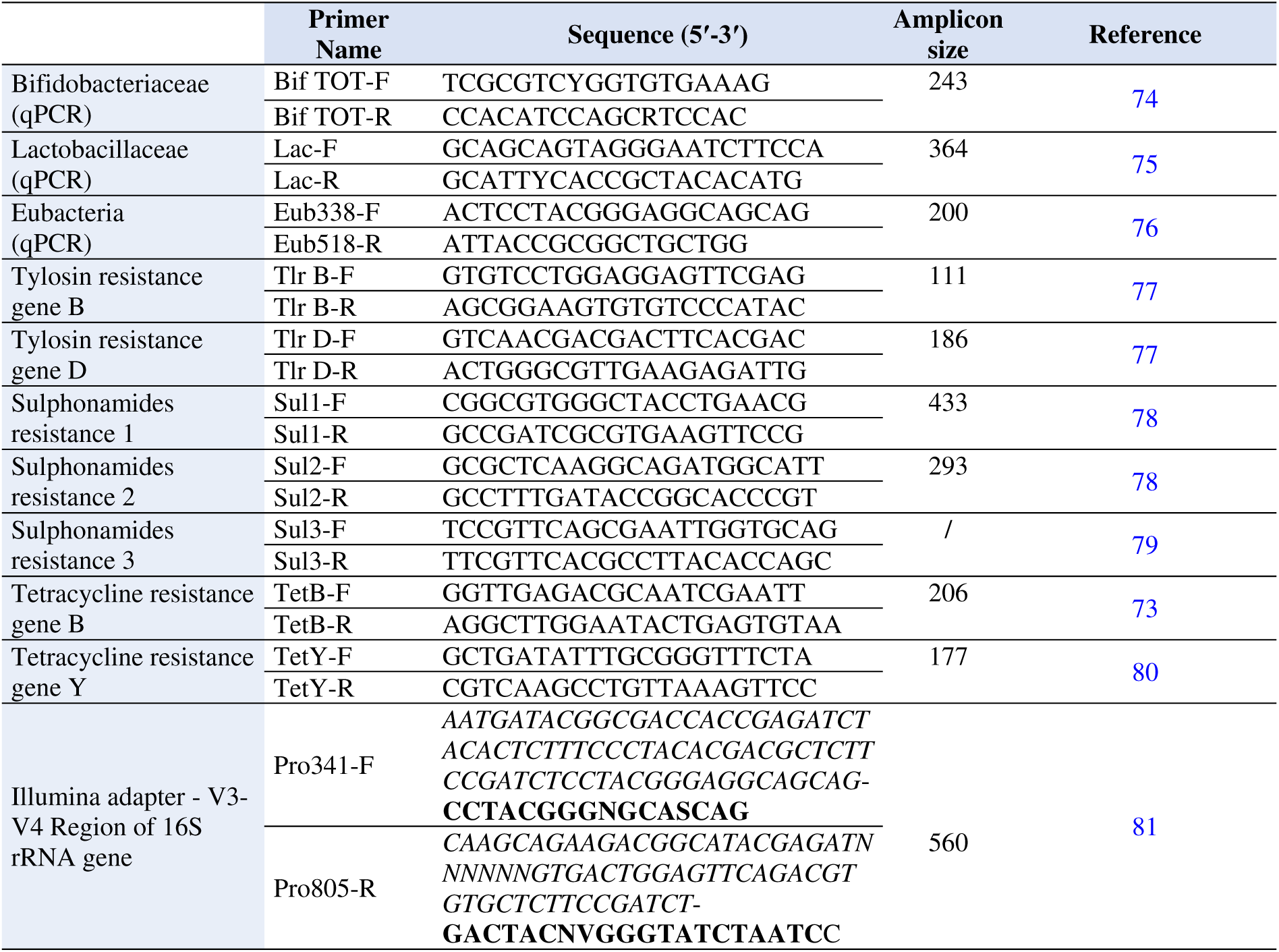
List of primers used in this experiment to carry out quantification of specific microbial targets, and detection of ARGs.

### Data adjustments and classification of microbial genera

Rarefied biom tables obtained from NGS bioinformatic analysis were used for further data adjustments: the absolute abundance of each bacteria species was calculated according to Raymann *et al*., (31), by multiplying absolute abundance data to the corresponding qPCR total amount results, and normalizing by the copy number of 16S rRNA gene typical of each microbial genus. Moreover, species belonging to the *Lactobacillus* genus have been recently re-classified (84) but databases for NGS OTUs assignment were not yet updated with the new classification at the time of the bioinformatic analysis of the presented data. Therefore, the absolute abundance dataset was manually curated to assign the former *Lactobacillus* spp. Firm-4 to *Bombilactobacillus* spp. genus and the former *Lactobacillus kunkeei* and *Lactobacillus plantarum* to the new respective taxonomical classifications *Apilactobacillus kunkeei* and *Lactoplantibacillus plantarum*. Due to the sequencing amplicon length (≃ 470 bp) might not be enough to efficiently discriminate among species, the manual curation was then validated in qPCR with Firm-4 and Firm-5 specific primers (33). The obtained dataset was used for further graphical and statistical analyses on target genera and species.

### Statistical analysis

Statistical analysis for qPCR and NGS data (α-diversity and taxon analysis) was performed with the R software (85) according to Baffoni *et al.* (68). Analysis on data normality and homoscedasticity was performed, therefore normal and homoscedastic data were analysed with ANOVA, non-normal, homoscedastic data (with normal distribution of residuals) were analysed with glm function, data with high deviation from normality where analysed with non-parametric Kruskal-Wallis test coupled with Dunn-test. For β–diversity index, data resulting from QIIME statistical elaboration were reported. The software calculates the UniFrac distance (weighted and unweighted UniFrac) between all the pairs of samples in the dataset to create a distance matrix. The statistical significance between groups was subsequently estimated using the Monte Carlo method with the Bonferroni correction. Post-hoc test among different groups was carried out and Bonferroni’s correction was applied. The post-hoc test considered pairwise comparisons within each experimental condition, taking into consideration the impact of each treatment over time. Therefore, four comparisons for the semi-field trial and three comparisons for the *in-field* trial were considered. The control was considered as a further treatment to monitor and evaluate the normal gut microbial community evolution resulting from the interaction of honeybees with the environment. Graphs were generated with ggplot2, ggpubr and Microsoft Excel. Biological relevance of experimental conditions, pairwise compared at their respective sampling time (T1 vs T0) was computed with Cramér’s V (86) relying on packages rcompanion, vcd, psych, desctools and epitools. Finally, PCA analysis was performed using packages FactoMineR (87) and factoextra (88), taking into consideration 71 taxa at species level.

## Data availability

These sequence data have been submitted to NCBI repository under the Sequence Read Archive (SRA) databases under accession numbers SAMN16442373-SAMN16442378; SAMN16442391-SAMN16442396; SAMN16442397-SAMN16442402; SAMN16442409-SAMN16442414; SAMN16442427-SAMN16442432; SAMN16442444-SAMN16442449; SAMN16442450-SAMN16442455 and SAMN16442462-SAMN16442467, Bio project n° PRJNA669646. Supplementary data, including exel files of elaborated data obtained from qPCR for target microbial groups and ARG and NGS data categorized at phyla, family and genera level, can be found at the Dryad Digital Repository (DOI…..).

## Compliance with ethical standards

This article does not contain any studies with human participants by any of the authors and experiments on animals were performed according to the Italian laws that allows experiments on arthropods without the need of an official ethical commission approval, unless cephalopods are used.

## Acknowledgments

The research was partially funded by the EU project “NOurishingPROBiotics to bees to Mitigate Stressors” (NO PROBleMS), H2020-MSCA-RISE 2017, GA 77760, 2018-2022. The founder had no role in the study design, data collection and interpretation, or the decision to submit the work for publication.

